# MALDI-TOF MS profiling and its contribution to mosquito-borne diseases: a systematic review

**DOI:** 10.1101/2024.07.18.604078

**Authors:** Monique Melo Costa, Vincent Corbel, Refka Ben Hamouda, Lionel Almeras

## Abstract

Mosquito-borne diseases are responsible for hundreds of thousands of deaths per year. The identification and control of the vectors that transmit pathogens to humans are crucial for disease prevention and management. Currently, morphological and molecular approaches are the standard methods used for vector identification, however, they present several limitations. In the last decade, matrix assisted laser desorption/ionization-time of flight mass spectrometry (MALDI-TOF MS) profiling emerged as an innovative technology in Biological Sciences and is now considered as a relevant tool for the identification of pathogens and arthropods. Beyond species identification, this tool can be relevant for the determination of different life traits of arthropod vectors. The purpose of the present systematic review was to highlight the contribution of MALDI-TOF MS to the surveillance and control of mosquito-borne diseases. Published articles from January 2003 to September 2023 were retrieved, considering different aspects of mosquito life traits which could be determinant in the transmission of diseases and vector management. The screening of scientific literature resulted in the selection of 54 published articles that assessed MALDI-TOF MS profiling to study various biological factors such species identification, life expectancy, gender, trophic preferences, microbiota and insecticide resistance. Although a large majority of the selected articles focused on species identification, the present review shows that MALDI-TOF MS profiling is promising for identifying various mosquito life traits with rapidity, high throughput capacity, reliability and low cost. The strength and weakness of this proteomic tool for vector control and surveillance are discussed.

## Introduction

Mosquitoes are insects that belong to the Culicidae family. At present, a total of 3,586 mosquito species have been identified worldwide, of which 88 are considered as important vectors of human diseases [1]. The primary mosquito vectors belong to the following genera: *Anopheles*, *Aedes* and *Culex* [1]. Mosquito-borne diseases (MBDs) have caused major outbreaks in human populations and account for about 700,000 deaths per year [2]. Malaria is responsible for more than half of the annual mortalities (n = 400,000), followed by dengue (n = 40,000) [2]. Mosquitoes are considered as the deadliest animals on Earth [3].

Tropical and subtropical areas are the most affected, but the intense worldwide transportation of people and goods and global warming have promoted the dispersion of invasive vectors in new areas [4,5]. In Europe, incursions of *Aedes albopictus*, one of the primary vectors of arboviruses, were reported in 26 countries [4,6–8]. This mosquito species was incriminated in local transmission of Chikungunya in Italy [9,10] as well as Chikungunya [11,12], dengue [13,14] and Zika [15] in France. The major dengue vector *Ae. aegypti* is also increasing its distribution worldwide and is now present in Southern Europe (Madeira Island, Turkey and Cyprus) because the conditions suitable for the vector to survive and prosper have expanded due to climate change [16]. The prevention of human-vector contact (e.g., nets, repellents) and the control of mosquito vector populations are the main strategies applied to limit MBD outbreaks. Chemical control with insecticides is the most used method to manage and prevent the spread of mosquitoes. However, the intense use of the few available compounds for decades has selected insecticide-resistant mosquito populations [17–20], reducing the efficiency of vector control based on chemicals.

Vector surveillance is a key component of all vector control programmes. Its main goals are to i) identify rapidly any changes in vector density, diversity, distribution, and insecticide resistance, ii) assess spatial and temporal risks of pathogen transmission and iii) guide timely decisions for vecotr control [21–23]. A rapid and accurate identification of vectors can prevent the establishment of invasive species in new territories by ensuring rapid and adequate vector control interventions [24]. Furthermore, the inventory of local mosquito fauna and its spatial-temporal evolution are of primordial importance for management programmes.

Morphological identification remains the conventional method for mosquito species classification [25]. It consists to research morphological structures of the specimen based on the use of the taxonomy keys [25]. Taxonomic identification is a cost-effective method and can be performed in the field. Although, it remains one the most widely used method for mosquito identification, morphological identification is time consuming and requires entomological expertise, which has been decreasing over the last 40 years [26]. In addition, specimen damages could occur during the sampling, transport or storing of mosquitoes. These subsequent alterations may conduct to incomplete species determination due to the loss of essential morphological criteria [27,28]. To circumvent the limitations of morphological identification, molecular techniques have been used as a relevant complementary approach. In general, the molecular method consists in the comparison of the nucleotide sequences of a molecular marker (i.e., DNA barcode), containing a unique genotypic feature for the analysed species, from a mosquito specimen with the known reference sequences, available in genomic databases (*e.g.*, GenBank, National Centre for Biotechnology Information [NCBI]; Barcode of Life Data Systems [BOLD] database). The confidence of the specimen identification at the species level is directly linked to the rate of matching sequence between the query and the database (*e.g.*, proportion of sequence homology/identity and scoring of accuracy) [29,30]. The mitochondrial cytochrome c oxidase subunit I (*COI*) is the commonly barcode gene used in mosquito identification, but additional genes, such the internal transcribed spacer (ITS2), are sometimes needed, notably for sibling or complex species. Despite its important contribution to a reliable mosquito identification and greater accessibility of this technology over time, molecular techniques remain relatively expensive and the specimens belonging to species complex are not always unambiguously identifiable [31,32]. The availability of molecular sequences from interest markers is another primordial factor to succeed in specimen identification.

In this context, matrix-assisted laser desorption/ionization time-of-flight mass spectrometry (MALDI-TOF MS) emerged as an alternative method for vector and pathogen identification. This technology was initially developed in the 1980s by Franz Hillenkamp and Michael Karas. They established a soft desorption ionization of particles using an organic compound called matrix, that gave rise to the name of the technique [33]. Then, the principle of MALDI-TOF MS consists in the ionization of sample particles mixed with a matrix through of a laser beam. The energy of the laser desorbs the matrix which transfers ions to sample molecules. These particles are accelerated in a tube under vacuum and the delay required by these particles to travel through of a tube until a detector, the time of flight (TOF), allows to determine their mass-to-charge ratio [33]. Finally, the detector transforms the energy measured for each ionized molecule into an electric signal and displays a spectral profile which mainly represents the most abundant, readily ionized small proteins and peptides with low masses (*i.e*. ranging generally from 2 to 20 kDa). The consideration of these spectra profiles as sample fingerprinting allows use of them for sample biotyping. The comparison of these protein profiles with a database of reference spectra created with known samples is indispensable for their classification [33]. Due to its simplicity, rapidity and high reliability, MALDI-TOF MS profiling has been used in routine for the identification of microorganisms since 2000’s [28].

The first articles reporting the use of MALDI-TOF MS for arthropod identification were published in 2005 and concerned the identification of fruit flies, *Drosophila melanogaster* [34] and aphid species [35]. Three years later, Dani and collaborators reported the differentiation of *An. gambiae* sensu stricto (*s.s*.) gender based on the comparison of their antennae’s MS profiles [36]. It took an additional five years to see the first report of successful identification of mosquitoes using MALDI-TOF MS; those works included the three major genera of the vectors, *Aedes* spp., *Anopheles* spp., and *Culex* spp. [37,38]. These studies achieved mosquito identification at the species level, even among specimens from *Anopheles gambiae* complex. Since then, this tool was successfully used to identify mosquitoes reared in the laboratory or caught in the field, confirming its efficiency to identify various mosquito species [39–42]. This success led the scientific community to extend the use of MALDI-TOF MS profiling for assessing other mosquito life traits such as the blood meal status [43–46], the population age and gender [47], the pathogen’s infection [48,49], the geographical origin [40,41,50] and the effect of mosquito sample storing and preparation [41,51–53]. More recently MALDI TOF MS was used to identify mosquito specimens throughout all development cycle (i.e., egg, larva, pupa, and adult stages) [37,38,54,55] and using different mosquito compartment (head, thorax, legs) to improve the sensitivity/accuracy of mosquito identification and/or to distinguish cryptic or close-related species [41,56,57]. Overall MALDI-TOF MS presents several advantages. The most significant one is the low cost of reagents, estimated at $1–2 per sample [40,58]. Another advantage is that MALDI TOF MS does not require specialized entomological skill and knowledge. Other advantages are the specificity per mosquito body part, the low volume of samples needed, and the fast sample processing. The main limitation to a wider application of MALDI-TOF MS profiling in medical entomology remains the high cost of the machine, around $200,000, and its maintenance [40]. Despite the high initial investment, once it is acquired by high throughput facilities, a fast return on investment is expected. Moreover, the wide application of this technology in microbiology laboratory diagnostic for routine identification of microorganisms such as bacteria, fungi and yeasts, contributed to its acquisition by numerous laboratories, excepted in low-middle income countries [28].

The aim of the present study is to review the current applications of MALDI-TOF MS profiling to analyse various mosquito life traits related to medical entomology and infectious diseases and then to underline the benefit that this tool may bring for the surveillance and control of mosquito-borne diseases.

### Methodology

The bibliographical search was performed following the Preferred Reporting Items for Systematic Reviews and Meta-Analyses (PRISMA) guidelines [59,60]. Scientific articles were retrieved from three publication databases: the Pubmed, the Web of Science and ScienceDirect. The searches were initially performed in February 2023 and updated manually in September 2023. The searches were carried out using specific search terms and their synonyms with the Boolean operator. In the Pubmed and the Web of Science, the search term was limited to “MALDI” AND “Mosquito”. The application of the same terms to ScienceDirect database allowed the retrieval of 5 times more articles (n = 676, with filters applied) than from the Pubmed and the Web of Science databases. The overview of article titles revealed that the majority was outside of the scope of the search terms “MALDI” and “Mosquito”, underlining that these terms were not adapted for ScienceDirect and that more specific terms should be used. The new search terms included “MALDI” or “mass spectrometry” associated with a term related to specific mosquito’s life traits (*e.g*. “MALDI” AND “mosquito identification”; “mass spectrometry” AND “mosquito identification”, etc.). The complete set of search terms with respective results of articles retrieved per publication databases is provided in the Additional file S1.

According to the PRISMA methodology, the initial process of screening for articles to be included in this study was performed as mentioned above. Additional filters were applied, including the publication year, language, and type of article. Subsequently, duplicate records were excluded manually, and the articles were screened based on its title and abstract to evaluate whether the subject relates specifically to the use of MALDI-TOF MS, mosquito and life traits, and/or mosquito control/monitoring/surveillance. The full text of the selected articles was retrieved for further analysis. Other relevant articles that were not retrieved from the databases or those which were published after the date of the initial search (*i.e*. between March and September 2023) were included manually to generate the most up-to-date review. The initial literature search was not limited in time, but since the technology related to MALDI-TOF MS profiling was developed from the 2000s, only peer-reviewed papers published in English from 2003 to February 2023 were selected from the databases. Review articles, conference proceedings, abstracts without the full text, case reports, case series, documents which were not peer-reviewed, and doctoral theses were excluded. Duplicates were likewise excluded. The assessment of risk of bias was performed by two of the authors of the present review through independent screening of the article titles and abstracts.

## Results

A total of 722 articles were initially selected from the repositories including 134 from Pubmed, 131 from Web of Science, and 457 from ScienceDirect. Among these articles, 200 were duplicate records and were then excluded. The application of exclusion criteria using filters in the search (the year of publication, language, conference proceedings, and abstracts only) resulted in the exclusion of 144 additional articles. Of the 378 remaining articles, 322 did not show pertinence with the subject using both title and abstract during the first screening and were then excluded.

The assessment of risk of bias was performed during the first screening. Two authors classified the articles using the following codes: not included (0), included (1), or maybe (2). In case of doubt or disagreement, a third author (senior researcher) intervened in the discussion. Of 378 articles retained during the first screening step, only 83 (*i.e.* 22 %) articles required a joint analysis of the authors to decide whether or not they would be included in the present review. The full text of the remaining 56 papers was retrieved, read, and analysed. During this second screening, six articles were excluded because they did not meet the inclusion criteria. In addition, four articles published after February 2023 and meeting the inclusion criteria were added manually, resulting in the final inclusion of 54 studies (i.e. 50 from database search and four added manually). The detailed flowchart illustrating the steps of article selection is presented in Figure 1.

**Figure 1.**
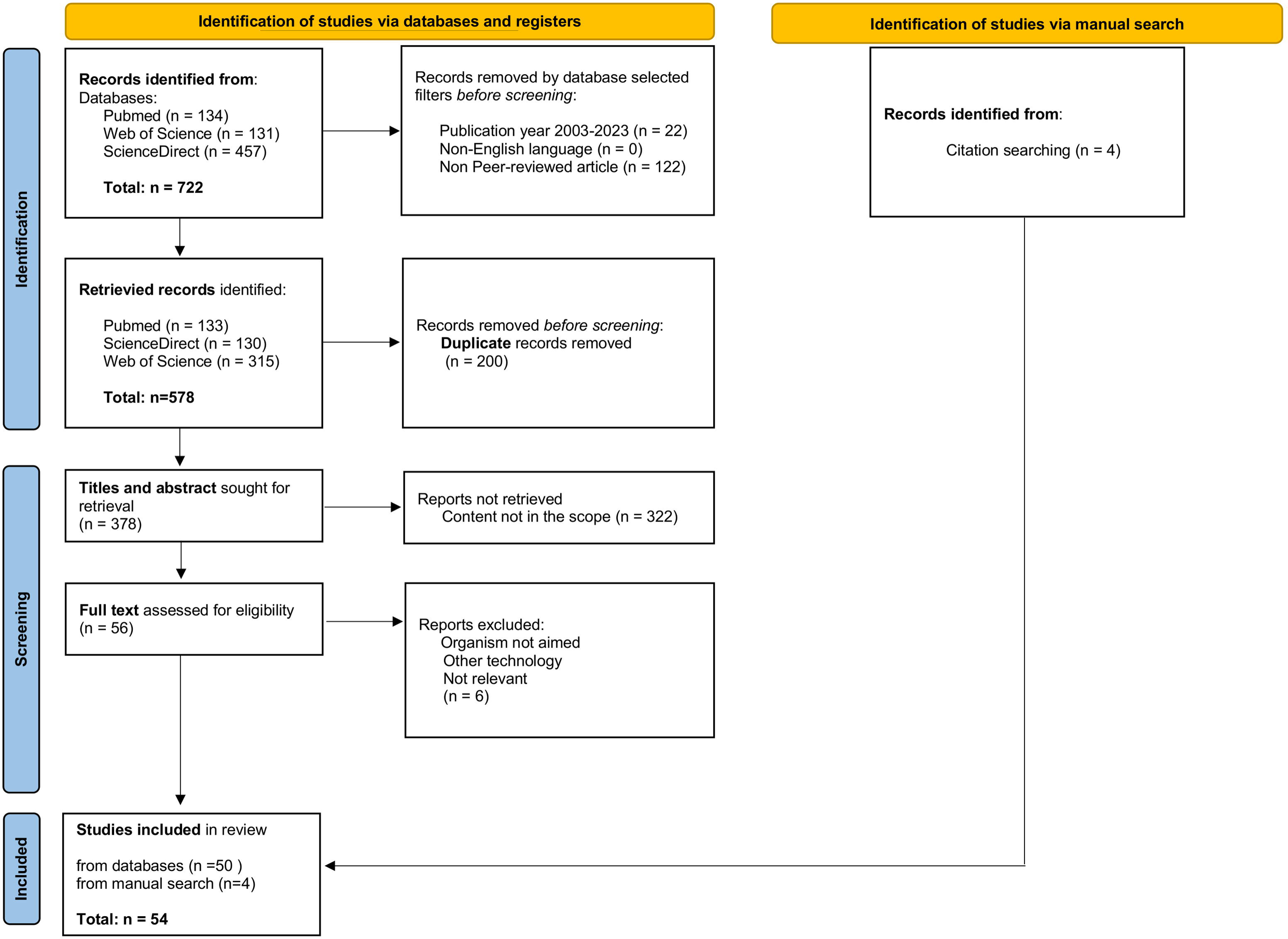
Flowchart showing selection process for a systematic review of the literature. Methodology and detailed results of the search, inclusion, and exclusion of studies followed the PRISMA model [108].

Among the mosquito life traits investigated, “mosquito identification” was the most common. The peak of articles relating to the use of MALDI TOF for mosquito analysis occurred in 2019 (Figure 2). France is the country that brought the largest contribution to the use of MALDI-TOF MS profiling for entomology studies in the last 20 years.

**Figure 2.**
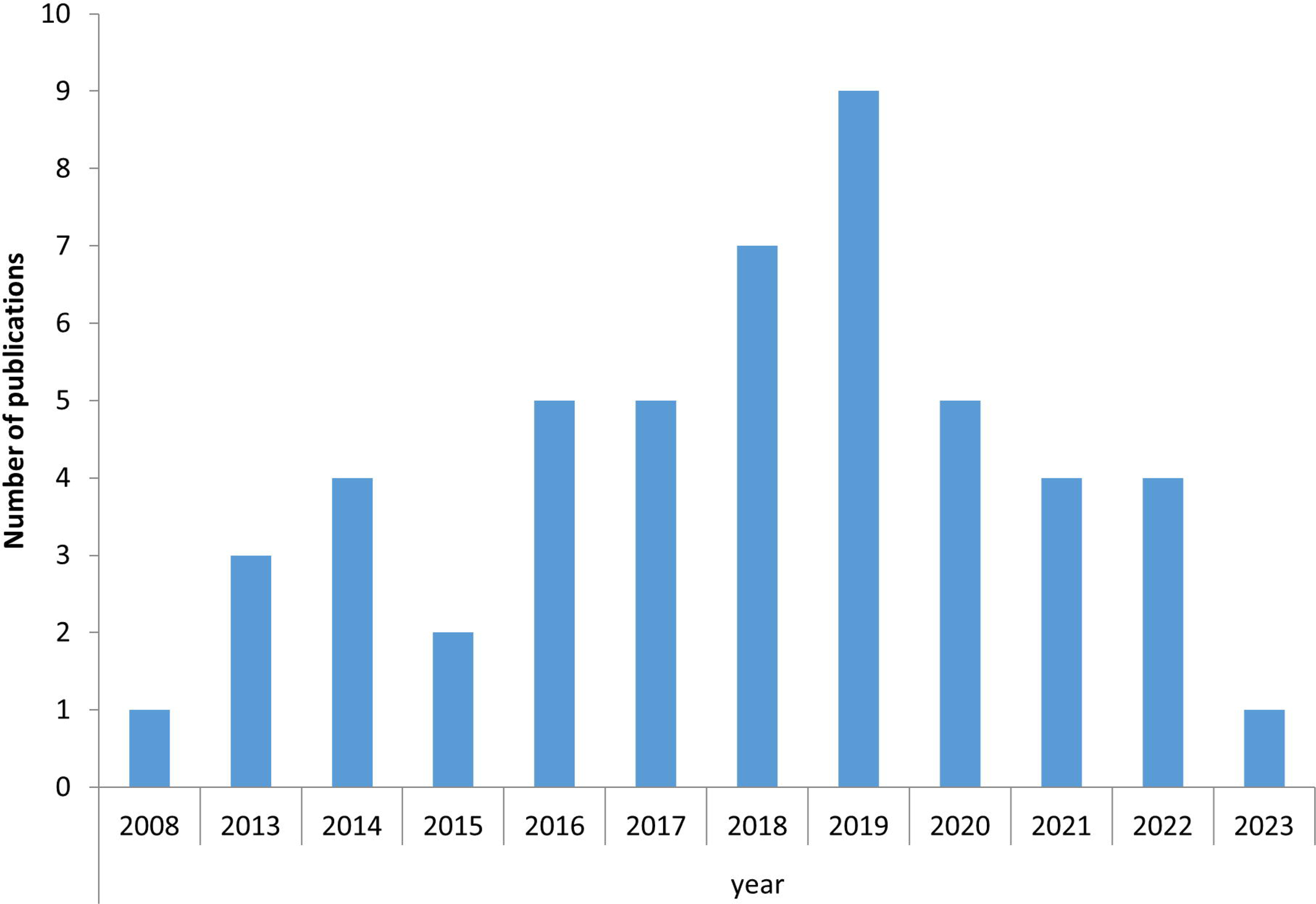
Number of publications reporting MALDI-TOF MS and mosquito life traits. Number of publications per year, from 2003 to 2023, retrieved from the scientific databases on MALDI-TOF MS profile and mosquito life traits.

The 54 articles included in the present study were then analysed and sorted according to the mosquito’s life trait analysed; i) species identification, ii) vector surveillance iii) identification of development stages, iv) mosquito life expectancy, v) tropic preference, vi) geographic origin, vii) pathogens infection viii) mosquito microbiota, and ix) mosquito resistance to insecticide. These sections are completed by a description of promising innovative applications of MALDI-TOF MS for entomological studies.

### 1. Assessment of MALDI-TOF MS profiling for species identification

Due to the complexity and time needed to correctly identify mosquito species using taxonomy method and DNA barcoding approaches, MALDI-TOF MS has emerged as a promising tool for systematic studies [25]. As indicated previously, the first application of MALDI-TOF MS profiling on mosquitoes occurred in 2008 and concerned the differentiation of *Anopheles gambiae* gender based on the comparison of MS spectra obtained from their antennae [36]. Since this pioneer work, other studies for mosquito identification were undertaken in 2013 [37,38]. These two studies performed independently demonstrated that MALDI-TOF MS could identify adult mosquitoes with high reliability. Müller and collaborators [37]then evaluated the capacity of MALDI-TOF MS to differentiate 12 distinct *Anopheles* species, essentially from laboratory origin, including members of the *An. gambiae* complex. MALDI-TOF MS achieved to differentiate these mosquito species with the spectra obtained from cephalothorax. Nevertheless, the separation of sibling species was incomplete based on hierarchical cluster analysis of the spectra. The authors failed to identify unique peaks which could be used as single biomarkers to distinguish these closely-related species. To overcome the limitations of unsupervised cluster analysis, a supervised statistical model was applied on these taxonomically closely related species. The application of this model on *An. gambiae* complex (*i.e*., *An. gambiae s.s.; An. arabiensis*; *An. merus* and *An. quadriannulatus*) demonstrated that 95% of laboratory-reared mosquitoes could be correctly classified. The application of the same model on M and S molecular forms of *An. gambiae s.s.* (the "M form" is now named *Anopheles coluzzii* while the "S form" retains the name *Anopheles gambiae* Giles), succeeded to classify correctly 91% of the specimens according to their molecular forms. This work demonstrated that MALDI-TOF MS could be used to discriminate anopheline mosquito species, even at the sibling-species level, with relatively high acuracy [37].

In the same year (*i.e*. 2013), Yssouf A. and colleagues assessed the performance of MALDI-TOF MS for mosquito identification using specimens from 20 species coming from 6 genera (4 *Aedes* spp., 9 *Anopheles* spp., 4 *Culex* spp., 1 *Lutzia* spp., 1 *Orthopodomyia* spp. and 1 *Mansonia* spp.) [38]. In contrast to the study conducted by Müller *et al*. [37], all the mosquito specimens were collected in the field and originated from La Reunion Island and Senegal. The authors selected legs as body part submitted to MALDI-TOF MS and obtained reproducible and high quality MS spectra among specimens from the same species. The introduction of leg MS spectra from each species or sibling-species in the reference database allowed them to correctly identify 100% of the specimens tested. The selection of legs for mosquito identification presents numerous advantages. The majority of the specimen body parts are preserved and could be used for other purposes, e.g. for the detection of pathogens in salivary glands or for the determination of parity rate through ovaries dissection. Finally, these first studies confirmed that MALDI-TOF-MS analysis of protein extracts from mosquito body parts was a suitable method for identifying different specimens until the distinction of sibling species [37,38]. These primary works also underlined that different body parts could be used for mosquito identification. As protein repertory changes according to body part for samples from the same species, the standardization of protocol is however required.

Effectively, the lack of guidelines and standardized protocols about the mosquito compartment selected for MS submission, including methods for sample preparation and homogenization can impair specimen identification. In the literature, legs were the mosquito part the most used for identification [38]. However, others studies demonstrated that different body part could be successfully used for imago identification, such as the head [37], cephalothorax [37,61], thorax or legs, either individually [38,50,62–65] or paired [53,56,57,66]. In 2016, Nebbak et al., proposed guidelines for body part selection, protocols of sample preparation, preservation mode, duration of storing and homogenization modes for mosquito identification at both larval and adult stages to facilitate MS spectra data exchange among laboratories [52]. For these demonstrations, two species from the *An. gambiae* complex (*An. coluzzii* and *An. gambiae s.s.*) were used. The reproducibility and stability of species-specific MS profiles were endpoints to define the optimized conditions. The authors concluded that the legs and whole larva appeared as the most adequate body parts for mosquito identification. They also showed that automatic homogenization methods of the samples were better than the manual one’s because they offer the possibility to test larger sample size without variation in homogenization performance in opposition with manual grinding. Moreover in the automatic modes, the addition of glass powder [52] or glass beads [56] as sample disruptor is recommended and an optimization of the homogenisation conditions are required according to the apparatus used [52]. Other studies comparing manual and automatic homogenization methods also highlighted the superiority of the automatic mode [41]. The automation and standardization of mosquito sample preparation methods for MALDI-TOF MS analyses overcome the time barrier posed by manual sample preparation and the problems of intra-species spectra heterogeneity.

To assess the storing mode and duration of storing, a kinetic submission to MALDI-TOF MS of mosquito samples stored under the usual conditions in the laboratory for different duration of time (one week to six months), revealed that freezing mode at -20°C or in liquid nitrogen (−196°C) for up to six months appears to be optimal for the identification of both development stages (adult and larval) [52]. Nevertheless, reliable identification was also obtained for adult mosquito specimens stored at room temperature with silica gel for up to 70 days. This method of preservation can be an alternative when freezing the samples is not possible, notably in the field. Conversely, the ethanol preservation mode was not recommended for testing mosquitoes in MALDI TOF MS [52].

Other study showed that the time that samples remain on a mosquito trap can accelerate the protein degradation and hence impact on the outcomes [51]. In addition, during the transport and storage of biological specimens, some parts of the mosquito can be damaged or lost due to handling, notably the legs which are breakables. It was repeatedly reported that the quality of the spectra was largely dependent on the number of legs submitted to MALDI-TOF MS and decreasing the the number of legs can impair the identification [51,52,67].

To circumvent the limitations of mosquito identification using legs, more recently other body parts were assessed. Among them, the thorax appeared as a relevant body part. It presents several advantages; it is not breakable and this relative large body part contains more proteins than legs and could then generated MS spectra of higher intensity [41]. To improve mosquito identification, the submission of two distinct body parts from the same specimen to MALDI-TOF MS was assessed [53]. As protein repertories are distinct per mosquito body part and as legs and thoraxes were already reported as relevant compartments for mosquito identification, it was proposed to submit independently to MS the two body parts (ie, legs and thorax) from a same specimen. The aim was to double-check the identification and to assess the concordance of identification between these two body parts. The application of paired samples submission to MS of mosquito field collected in Guadeloupe Island revealed concordant species identification (100%) in agreement with morphological classification [53]. This double-checking system is particularly relevant when close-related species have to be identified or when the quality of the MS spectra from one body compartment is insufficient (eg, loss of several legs). This strategy was applied for the distinction of 8 *Anopheles* spp. [56] and 13 *Culex* spp. [57], including species which could not be distinguished morphologically. A concordant, correct and relevant identification was obtained for all spectra from both body parts of these last two studies. Interestingly, one specimen classified as *An. peryassui* using taxonomic criteria was identified as *An. intermedius* based on MALDI-TOF MS analysis of legs and thoraxes [56]. The confirmation of MS results by molecular biology underlined the accuracy of the proteomic classification. It is noteworthy that the paired submission to MS of double body part per specimen improved the identification rate by increasing the associate identification score and the concordance of the results. This paired MS query may be decisive for the distinction of cryptic species.

Although head, thorax, and legs are suitable for mosquito identification with MALDI-TOF MS, the performance of the identification can vary for a same specimen depending on the body part used [40,41]. Nabet et al. contested that thoraxes from mosquitoes was the better body part for specimen identification by MS, claiming that better results were obtained with mosquito head [40]. These authors reported that one limitation of the use of thorax was the risk of spectra alteration in freshly engorged mosquitoes due to the presence of blood trace in this body part, as observed with some field collected specimens [40]. In another study, the comparison of MS spectra from different body parts of artificially blood-fed mosquitoes confirmed the detection of characteristic MS peaks of blood in the thorax [41]. However, the same MS peaks indicating the presence of trace amounts of blood were also detected in the spectra generated by the legs and head of the same specimens. These peaks in mosquitoes engorged with human blood were located at 15 138 m/z and 7568 m/z of MS profiles [41]. These both MS peaks were already observed in MS spectra profiles from human blood [43]. The blood traces were then attributed essentially to contamination of safe tissues which should occurred during the specimen dissection of engorged mosquitoes. The absence of these blood-associated MS peaks in the spectra of several thorax samples from engorged specimens, as well as the much less frequent detection in leg samples, supports this hypothesis. It is interesting to note that the blood-associated peaks detected in MS spectra from different body parts tested did not hamper specimen identification. Nevertheless, this blood contamination could decrease the identification score. According to the authors, the order of preference of the body parts to be analysed to obtain an optimum spectral profile for mosquito identification is as follows: thorax, legs, and head [41]. These findings are in agreement with other recent studies published in this field [53,56,57].

Nowadays, the main limitation for a wide-scale use of this proteomic tool for the identification of mosquitoes and their life traits is the absence of a central, free acces, reference database of MS spectra. To overcome this limitation, it is essential that investigators of each study share their reference spectra database [56,57]. The procurement of a large diversity of mosquito species from distinct origins and developmental stages is then required to enrich reference spectra DB. At present, MS profiles for at least nine genera of mosquitoes, encompassing 67 species, are available. Of these nine genera, the highest numbers of MS profiles have been characterized for mosquitoes from the *Anopheles*, *Culex,* and *Aedes* genera. The other six genera consist of a single species (Table 1). The MS databases include different mosquito developmental stages, from eggs to adult, which further highlights the potential of MALDI-TOF MS for studies on mosquitoes.

**Table 1.** Availability of spectral profiles for mosquito species in reference MS database.

| Genus (n=9) | Species (n=67) | Immature | Imago (compartment) | Reference* |
| --- | --- | --- | --- | --- |
| *Anopheles* | *An. aquasalis* |  | Adult (Legs and Thorax) | [56] |
|  | *An. braziliensis* |  | Adult (Legs and Thorax) | [56] |
|  | *An. darlingi* |  | Adult (Legs and Thorax) | [56] |
|  | *An. nuneztovari* (s.l.) |  | Adult (Legs and Thorax) | [56] |
|  | *An. triannulatus* (s.l.) |  | Adult (Legs and Thorax) | [56] |
|  | *An. oswaldoi* (s.l.) |  | Adult (Legs and Thorax) | [56] |
|  | *An. intermedius* |  | Adult (Legs and Thorax) | [56] |
|  | *An. peryassui* |  | Adult (Legs and Thorax) | [56] |
|  | *An. gambiae* | Larvae, Pupae | Adult (Head, Thorax and Legs) | [38,40,41,52,71] |
|  | *An. coluzzii* | Larvae, Pupae | Adult (Legs and Thorax) | [41,52,71] |
|  | *An. funestus* |  | Adult (Head, Thorax and Legs) | [38] |
|  | *An. ziemanni* |  | Adult (Legs) | [38] |
|  | *An. arabiensis* |  | Adult (Head, Thorax and Legs) | [38,40] |
|  | *An. wellcomei* |  | Adult (Legs) | [38] |
|  | *An. rufipes* |  | Adult (Legs) | [38] |
|  | *An. pharoensis* |  | Adult (Legs) | [38] |
|  | *An. maculipennis* | Larvae |  | [68] |
|  | *An. claviger* |  | Adult (Legs) | [62] |
|  | *An. hyrcanus* |  | Adult (Legs) | [62] |
|  | *An. maculipennis* |  | Adult (Legs) | [62] |
|  | *An. bancroftii* |  | Adult (Legs) | [51] |
| *Culex* | *Cx. dunni* |  | Adult (Legs and Thorax) | [57] |
|  | *Cx. nigripalpus* |  | Adult (Legs and Thorax) | [57] |
|  | *Cx. quinquefasciatus* |  | Adult (Legs and Thorax) | [38,51,53] |
|  | *Cx. usquatus* |  | Adult (Legs and Thorax) | [57] |
|  | *Cx. adamesi* |  | Adult (Legs and Thorax) | [57] |
|  | *Cx. dunni* |  | Adult (Legs and Thorax) | [57] |
|  | *Cx. eastor* |  | Adult (Legs and Thorax) | [57] |
|  | *Cx. idottus* |  | Adult (Legs and Thorax) | [57] |
|  | *Cx. pedroi* |  | Adult (Legs and Thorax) | [57] |
|  | *Cx. phlogistus* |  | Adult (Legs and Thorax) | [57] |
|  | *Cx. portesi* |  | Adult (Legs and Thorax) | [57] |
|  | *Cx. rabanicolus* |  | Adult (Legs and Thorax) | [57] |
|  | *Cx. spissipes* |  | Adult (Legs and Thorax) | [57] |
|  | *Cx. nigripalpus* |  | Adult (Legs and Thorax) | [53] |
|  | *Cx. pipiens* | Larvae, Pupae | Adult (Legs) | [38,62,68,71] |
|  | *Cx. modestus* |  | Adult (Legs) | [62] |
|  | *Cx. hortensis* | Larvae |  | [68] |
|  | *Cx. atratus* (s.l.) |  | Adult (Legs and Thorax) | [53] |
|  | *Cx. iyengari* |  | Adult (Legs) | [51] |
|  | *Cx. sitiens* |  | Adult (Legs) | [51] |
|  | *Cx. annulirostris* |  | Adult (Legs) | [51] |
|  | *Cx. molestus* | Larvae and Pupae |  | [71] |
| *Aedes* | *Ae. aegypti* | Eggs, Larvae, Pupae and Exuviae of larvae and pupae | Adult (Legs and Thorax) | [38,41,51,53,55,71,72] |
|  | *Ae. albopictus* | Eggs, Larvae, Pupae and Exuviae of larvae and pupae | Adult (Legs and Thorax) | [38,41,52,53,55,68,71,72] |
|  | *Ae. atropalpus* | Eggs |  | [55] |
|  | *Ae. cretinus* | Eggs |  | [55] |
|  | *Ae. geniculatus* | Eggs |  | [55] |
|  | *Ae. japonicus* | Eggs |  | [55] |
|  | *Ae. koreicus* | Eggs |  | [55] |
|  | *Ae. phoeniciae* | Eggs |  | [55] |
|  | *Ae. triseriatus* | Eggs |  | [55] |
|  | *Ae. taeniorhynchus* |  | Adult (Legs and Thorax) | [53] |
|  | *Ae. cinereus* |  | Adult (Legs) | [62] |
|  | *Ae. vexans* |  | Adult (Legs) | [51,62] |
|  | *Ae. caspius* |  | Adult (Legs) | [62] |
|  | *Ae. rusticus* |  | Adult (Legs) | [62] |
|  | *Ae. excrucians* |  | Adult (Legs) | [62] |
|  | *Ae. scutellaris* |  | Adult (Legs) | [51] |
|  | *Ae. notoscriptus* |  | Adult (Legs) | [51] |
|  | *Ae. vigilax* |  | Adult (Legs) | [51] |
| *Mansonia* | *M. uniformis* |  | Adults (Legs) | [38] |
| *Culiseta* | *Cs. longiareolata* | Larvae |  | [68] |
| *Ochlerotatus* | *Oc. caspius* | Larvae |  | [68] |
| *Deinocerites* | *D. magnus* |  | Adult (Legs and Thorax) | [53] |
| *Psorophora* | *P. cingulata* |  | Adult (Legs and Thorax) | [53] |
| *Coquillettidia* | *Cq. richiardii* |  | Adult (Legs) | [62] |

\*All the mosquito species identification was done by MALDI-TOF MS with a Microflex LT MALDI-TOF Mass Spectrometer (Bruker Daltonics), at the exception of the work realized by Schaffner et al [55] that used MALDI-TOF Mass Spectrometry Axima™ Confidence machine (Shimadzu-Biotech).

### 2. Assessment of MALDI-TOF MS profiling for mosquito surveillance

The successful use of MALDI-TOF MS profiling for the identification of mosquitoes led to its application in surveillance programmes. In the present review, four studies that applied this novel technology from medium to large scale monitoring programmes were found [6,24,58,68]. MALDI-TOF MS was selected in these studies due to its ease, rapidity, and low cost in identifying mosquito species, compared to the traditional method based on morphological identification or DNA barcoding. Although the transmission of pathogenic agents by mosquitoes occurs mainly during the adult stage, the monitoring and control were generally carried out during the pre-immature (egg) or immature (larvae) stages. Among the four studies that applied MALDI-TOF MS for mosquito surveillance, three relied on egg colection whereas one relied on larval stages.

The studies using mosquito eggs were conducted in Swiss-Italian border [24,58]and the United Kingdom [6]. In the British study, MALDI-TOF MS contributed, for the first time to the successful identification of the invasive *Ae. albopictus* [6]. In addition to morphological identification, scanning electron microscopy and direct sequencing of PCR products of *COI* and *ITS2* genes confirmed the identification of the eggs obtained by MALDI-TOF MS. This work revealed the robustness of the proteomic tool to monitor the incursions of *Ae. albopictus* in the United Kingdom.

In southern Switzerland, a surveillance programme on *Ae. albopictus* was implemented in 2000 for 13 years [24]. This survey was carried out because the neighbouring areas on the Italian side of the border were considered as high risk for the introduction of *Ae. albopictus*. *Ae. albopictus* was identified for the first time in 2003 in Switzerland in the canton of Ticino, located along the Swiss-Italian border. Initially, the eggs, larvae, and adult mosquitoes were submitted to taxonomic identification. MALDI-TOF MS was subsequently used to confirm the identification of the eggs, given that the classical morphological identification of mosquitoes collected with ovitraps was considered as unrealistic for such a long entomology suvery. MALDI-TOF MS succeeded in distinguishing the eggs of *Ae. albopictus* from those of *Ae. geniculatus,* despite the fact that these two *Aedes* mosquitoes are in sympatry and their eggs are not easily distinguishable morphologically [24]. More than 3,000 eggs analysed by MALDI-TOF MS were identified as *Ae. albopictus*. The timely surveillance measures implemented along the Swiss-Italian frontier helped to limit the introduction and spread of this vector in this territory hence reducing the risk of arbovirus tranmssion.

As part of the Swiss-Italian border monitoring programme [58], MALDI-TOF MS was also used to identify the introduction of another invasive species, *Aedes koreicus*. Native from Asia, *Ae. koreicus* is competent to transmit the Japanese encephalitis virus and dog heartworm *Dirofilaria immitis*. Identified for the first time in 2013 by MALDI-TOF MS, this invasive mosquito species is spreading in Central Europe [58]. A validated MS database, curated at Mabritec SA (Riehen, Switzerland) was used as the reference for mosquito identification. Given the low hatching rate, morphological identification of larvae and adults was unfeasible. MALDI-TOF MS was therefore applied as a more practical alternative to distinguish the new invasive *Ae. koreicus* from *Ae. albopictus*. This surveillance programme showed that, in case of a low hatching rate of an invasive mosquito, MALDI-TOF MS can be pertinent to rapidly identify invasive species and could then replace the challenging and laborious procedure of morphological identification of mosquito larvae.

Finally, in the south of France, MALDI-TOF MS technology was applied during the summer of 2015 to monitor the presence of mosquito larvae in an urban area of Marseille [68]. Among 2,418 larvae or pupae submitted to MALDI-TOF, 93.4% (n= 2,259) were correctly identified, hence distinguishing five species. The lower rate of relevant identification was obtained for early instar larvae and/or pupae. The lower protein concentration of the early instar larvae and metamorphosis which occurs at the pupal stage are likely to explain the lower matching rate of MS spectra with those of the references. Interestingly, *Culex impudicus*, a mosquito species which was not initially included in the database, was nevertheless detected. This result confirmed the high species-specificity of MS spectra. The combination of the results of species abundance with twelve physicochemical variables of larval habitats allowed to shed light on the association between specific environmental factors and the presence of mosquito species. Collectively, these works confirmed the usefulness of MALDI-TOF MS for large-scale monitoring of mosquitoes at immature stages and may contribute to improve the efficiency of mosquito control programmes.

### 3. Assessment of MALDI-TOF MS profiling to mosquito identification according to the developmental stages

Numerous studies have demonstrated the successful application of MALDI-TOF MS profiling for identification of adult mosquitoes using different body parts [25,28,69,70]. The only body part of adult stage which is not recommended for mosquito identification by MALDI-TOF MS is the abdomen [25]. The principal reason is the presence of mammalian host blood and/or digested food in the digestive tract which produce extraneous or artefactual MS spectra. The latter spectra associated with the presence of food or host blood usually render the acquisition of species-specific protein signature impossible.

Some authors also demonstrated that the application of MALDI-TOF MS for mosquito identification can be extended to the immature stages (*i.e*., egg, larva, and pupa) [55,58,68,71]. Although the lower protein concentration present in the early instars of larvae (L1/L2) may not allow an accurate specimen identification by MS [54,68], when late instars of larvae (L3/L4) are used, the rate of correct and relevant identification exceeds 92%, which is largely sufficient for surveillance programmes. For the identification of larvae, the use of the whole body is recommended because the dissection of larvae could be laborious and time-consuming, notably for early stages, and also because incomplete dissection may lead to spectral heterogeneity [71]). Since the gut content could alter intra-species reproducibility of MS spectra, this parameter was assessed for whole larvae. Different diets given to laboratory-reared mosquito larvae have a minor effect on MS profile patterns, confirming the suitability of the use of the entire specimen at immature stages [68].

As for the eggs in the context of monitoring programmes, the ovitraps used for *Aedes* monitoring are frequently laden with eggs, and the identification of the eggs at the same time as the adult stage may improve the efficacy of vector surveillance. Schaffner *et al*. [55] tested whether it was possible to identify a pooled mixture of eggs from different *Aedes* species. Eggs from the same species and pools of ten aedine eggs including two or three distinct *Aedes* species in different ratios were assessed by MALDI-TOF. All nine aedine species in a collection of eggs from a single species, as well as two or three *Aedes* species in mixed pools of ten eggs were identified correctly by MALDI-TOF MS. The minimum of three eggs per species was necessary in the pools for the identification of the species [55].

One of the problems of mosquito identification by MALDI-TOF MS is the requirement for euthanasia of the specimens for MS submission, whatever the life stage. The euthanasia of the specimen impedes the performance of other assays requiring live mosquitoes, such as the evaluation of insecticide susceptibility. Seen from this point of view, analysis of mosquito exuviae (*i.e*., the outer skin that is shed off after a moult during the aquatic stages) appeared as an alternative. Nebbak *et al*. established the proof of concept of the application of MALDI-TOF MS for species identification using exuviae from the fourth-instar and pupal stages of laboratory-reared *Ae. albopictus* and *Ae. aegypti* [72]. The exuviae of each of these two aquatic stages yielded a distinct and reproducible MS spectral profile. All exuviae samples correctly identified the species. Despite the successful use of this shedded body part, few disadvantages were found. The cuticles of exuviae are sturdy and this feature limits protein extraction from exuviae. The resulting MS spectra then possessed a low peak diversity. The quick degradation of the exuviae in the field decreases protein abundance leading to identification impairment of resulting MS spectra. The MS analysis of exuviaes was not adapted to determine mosquito biodiversity history of larval habitat. On the other hand, the use of exuviae offers some advantages, including double confirmation of species identification using both fourth-instar and pupal exuviae from the same specimens, and the preservation of live material for other work.

The preceding sections highlighted that MALDI-TOF MS biotyping is suitable for mosquito identification at different life stages (eggs, larvae, pupae, adult and exuviae) and using different body compartments, and can then effectively contribute to vector surveillance. It was emphasized that it is necessary to follow standardized procedures, from the choice of samples to the methods of sample handling and treatment. These are the key points to obtain a reproducible specific MS protein profile. The addition of species-specific internal biomarker mass sets as calibrators in the samples, as previously suggested [50,55], considerably improves the performance of this proteomic tool.

### 4. Assessment of MALDI-TOF MS profiling to determine mosquito life expectancy

The transmission of mosquito-borne diseases is directly associated with vector competence and capacity. Vector competence is the ability of the vector to become infected and transmit pathogens. Vector capacity is the set of components of a vector population that determines its potential to transmit pathogens. The probability of daily survival and extrinsic incubation period (*i.e*., the period of pathogen multiplication in the salivary glands of the mosquito after an infected blood meal) comprise these components. In this sense, mosquito life expectancy is a key parameter influencing the risk of disease transmission. Mosquitoes that survive longer, have a higher probability to be infected. For example, to transmit malaria parasites, *An. gambiae* females must be older than 10 days to become infective [73]. The more often the female feeds on humans’, the higher is the probability to get infected by malaria parasites. Traditional methods to measure the mosquito age include the dissection of ovaries to assess the parous rates, analysis of cuticular lipid profiles by gas chromatography–mass spectrometry [74,75], near-infrared spectroscopy and, more recently, the measurement of the transcription levels of aging marker genes [76,77]. These techniques are labour-intensive, time-consuming and are currently not adapted to high throughput analysis. In this context, MALDI-TOF MS represents an interesting alternative for age grading studies.

MALDI-TOF MS was used for the first time to measure the age of mosquito using spectra derived from cuticular lipids [73]. The findings revealed that adult female *An. gambiae* mosquitoes old enough to transmit malaria parasites have different spectral profiles than the younger females. The MS spectra of cuticular lipid of female adults aged of one day, seven to ten days, and 14 days can be clearly distinguished by MALDI-TOF MS procedures. Mated females aged between seven and ten-days presented a three-fold increase in signal intensity at 570 m/z and 655–660 m/z compared to one-day-old virgin females, but the 570 m/z signal was suggested to belong to the mating signal. In contrast, a decrease in the signal intensity at 535–545 m/z was noted with increasing age. Higher abundance of lipids at 670–680 m/z and 700–710 m/z was found in 14 days old virgin females. An accurate analysis of MS spectra with multivariate statistical methods revealed that cuticular lipid profiles could be effective to detect differences between males and females or between virgin and mated females. MALDI-TOF MS proved to be efficient in determining the age of adult mosquitoes through the intensity and diversity of cuticular peaks. This tool can therefore contribute to risk assessment in vector control programmes, in particular for estimating the life expectancy of adult mosquitoes before and after implementation of vector control interventions.

More recently, Piarroux *et al*. evaluated the capacity of MALDI-TOF MS to classify the spectral patterns of laboratory-reared *An. stephensi* mosquitoes based on entomological drivers of malaria transmission, notably the age of specimens [78]. To detect protein fingerprint associated with mosquito age (0–10 days, 11–20 days and 21–28 days), MALDI-TOF MS was coupled with machine-learning algorithms (artificial neural networks, ANNs). This sophisticated bioinformatics analysis successfully detected specific *Anopheles* protein profile changes, allowing an association of spectral patterns with mosquito age, with an accurate prediction rate of 73%. The best results were obtained with MS spectra from the thorax. The predicted age groups were not associated with specific spectral peaks, but rather with the variations in peak intensity. More recently, the same team improved the age prediction reliability of *Anopheles* mosquitoes by optimizing deep learning frameworks used for MALDI-TOF MS spectra analyses [79]. The application of MALDI-TOF MS tool combined with machine learning approach to malaria vectors collected in the field succeeded to estimate mosquito age with an error of less than 2 days in the best conditions.

These three studies underlined that the estimation of mosquito age by MALDI-TOF MS profiling requires complex analyses and it should be done by analysing either cuticular lipid profiles [73] or protein patterns of the thorax [78]. The prediction of mosquito age, notably in human malaria vectors, remains essential to evaluate the risk of disease transmission and deploy adequate vector control interventions.

### 5. Assessment of MALDI-TOF MS profiling to determine mosquito trophic preferences

The determination of blood meal sources in mosquito vectors is essential to improve knowledge about host-vector interactions and pathogen’s transmission risk. The traditional methods of identification of the mammalian source of blood meal include serological tests, such as precipitin and enzyme-linked immunosorbent assay (ELISA), and molecular tests [44,80,81]. The major limitations of serological tests are unavailability of antibodies against a large range of potential hosts and specificity of antibodies [44]. Despite the successful application of molecular methods to detect the host blood-meal source, there are several limitations to this approach. The high cost of molecular assays, the poor quality of blood sample that may have undergone digestion in the engorged mosquito gut, the difficulties in analysing mixed blood meal sources [44,82], or the reliance on the availability of complete DNA sequence in public databases [46], are striking examples.

The first work aiming to assess the relevance of MALDI-TOF MS for blood meal identification was performed on laboratory-reared *An. gambiae* mosquitoes that were artificially engorged on seven distinct blood sources from vertebrates (human, horse, sheep, rabbit, mouse, rat, and dog) [43]. The mosquito abdomen samples containing blood proteins were kinetically submitted to MALDI-TOF MS. Specific MS profiles from engorged mosquitoes were observed according to host blood source, independently of the mosquito species. Kinetic analysis revealed that abdominal protein spectra remained stable up to 24 h post-feeding. After this time point, digestion of blood proteins induced alteration of MS profiles, reducing the proportion of correct identification of blood source, as with analysis using molecular biology success [43].

Since this primary work, two further studies enlarged the reference MS spectra database by testing the blood of 18 additional vertebrates, hence confirming the specificity of abdomen MS profiles from freshly engorged mosquitoes (*i.e*., ≤ 24 h) [44,63]. Interestingly, blood samples from three primates (*i.e*., *Callithrix pygmaea* [pygmy marmoset], *Erythrocebus patas* [hussar monkey], and *Papio hamadryas* [Hamadryas baboon]) were tested, and no mismatch with human blood occurred, supporting the high specificity of the spectra. Actually, blood samples from a total of 25 distinct hosts have been tested, and their MS spectral profiles deposited in the database. It was also demonstrated that mixed blood meals from mosquitoes which fed on two distinct hosts were also correctly identified by MALDI-TOF MS [46]. The high capacity of this proteomic tool to identify the blood source is important for identification of vectors responsible for the transmission of zoonotic pathogens and determination of animal reservoirs. In the field, it is generally recommended to crush mosquito abdomens on Whatman filter papers to stop blood digestion, MALDI-TOF MS analysis of dried blood samples on Whatman filter papers showed successful to identify the origin of blood meal [45]. MALDI-TOF MS succeeded to identify mosquito species and their respective host for blood fed specimens collected in five ecological areas of Mali [63]. The rate of successful classification reached nearly 93% (n=651/701) using Whatman filter.

More recently, the combination of machine-learning algorithms with MALDI-TOF MS approach allowed to distinct past-engorged mosquitoes from unfed mosquitoes by analysing mosquito thorax or legs with a success rate about 80% [78]. In this study, the origin of blood sources was not investigated for, since the aim was to distinguish between females that laid eggs (*i.e.*, parous specimens) from those that have not (*i.e.*, nulliparous), to estimate the risk of disease transmission.

These studies reinforce the relevance of MALDI-TOF MS to determine the feeding patterns of freshly blood fed mosquitoes. However, further experiments are required to enlarge the database of the mammalian sources of blood fed mosquitoes. The determination of blood meal sources is important to improve our knowledge on human-vector interactions and to identify reservoir hosts.

### 6. Impact of the geographic origin of mosquitoes on the MALDI-TOF MS profiling

A reliable and effective entomological monitoring programme requires a comprehensive database of reference spectra that covers relevant mosquito species involved in human pathogen transmission [62]. The reliability of MS identification is linked not only to the level of spectra specificity, but also to spectra reproducibility. Several factors could impair the generation of MS spectra, as presented in earlier sections. There are extrinsic factors, such as the method of sample preparation and storage, and intrinsic ones such as blood contamination, environmental conditions (*e.g*., climate, nutrition, microbial exposure, and population size) and genetic background of the specimen collected. Some studies have reported that variations in MS spectra can occur in larval and adult specimens belonging to the same species, proceed in the same conditions, but collected in distinct geographic areas [50,62,71]. These intra-species variations were attributed to changes in protein content which could be influenced by environmental factors and/or genetic background [71].

The variations of MS spectra among specimens from distinct geographic origins did not hamper species identification [41,50], with very few exceptions [40]. A study reported that the inclusion of three cosmopolitan mosquito species (*i.e*., *Ae. aegypti*, *Ae. albopictus*, and *Cx. quinquefasciatus*) in the database of mosquitoes from the Pacific region was sufficient to identify mosquito species from Asia and Africa with relevant scores [5]. The low rate of misidentification (2.0%) further indicated that MALDI-TOF MS is a robust method for mosquito identification at a global scale. Nevertheless, the introduction of specimens from the same geographic areas in the database significantly improved the accuracy of identification scores [40,41]. The available evidence suggests that the creation of region-specific reference MS databases could provide higher confidence scores and then improve the quality of mosquito identification.

### 7. Assessment of MALDI-TOF MS profiling to detect pathogens in mosquitoes

Detection of pathogens in mosquitoes is important to identify hotspots of disease transmission risk but also to estimate the intensity of transmission. An early detection of pathogens in the vector can lead to a timely and cost-effective responses and hence prevent outbreaks. Few works have investigated the efficiency of MALDI-TOF MS to distinguish infected versus non-infected mosquitoes. A limited number of published studies have focused on the detection of parasitic agents, including three filarioid helminths: *Dirofilaria immitis* (dog heartworm), *Brugia malayi* (Malayan lymphatic filarial worm, an etiologic agent of lymphatic filariasis), and *Brugia pahangi* (lymphatic filarial worm, an etiologic agent of lymphatic filariasis) in *Ae. aegypti* [48]; and malaria parasites (*Plasmodium berghei*) in experimentally-infected *An. stephensi* mosquitoes [49,78]. To avoid bias due to environmental factors, these studies were performed on laboratory-reared and artificially infected mosquitoes. In each of these studies, several mosquito body parts were tested. The performance of MS classification was assessed by comparing the results from MALDI-TOF MS analysis with the molecular analysis, the last method being the gold standard. The results showed that the highest performance, *i.e*., high sensitivity (86.6%, 71.4%, and 68.7% for *D. immitis*, *B. malayi*, and *B. pahangi*, respectively) and high specificity (94.1%), were obtained using the cephalothoraxes [48]. Among 37 MS peaks that discriminated between uninfected and infected *Ae. aegypti*, two MS peaks (4073 and 8847 Da) were specific to *Ae. aegypti* infected with microfilariae regardless of the nematode species or mosquito compartment. These two peaks were then considered as biomarkers for *Ae. aegypti* infected with these microfilariae.

In 2017, Laroche *et al*. tested the efficacy of MALDI-TOF MS in detecting changes in the protein profiles of *An. stephensi* experimentally infected with *P. berghei* [49]. The MS screening of uninfected and infected *An. stephensi* mosquitoes revealed concordant results (98.8%) with those of molecular methods. The differences between uninfected and infected groups of mosquitoes were attributed to variation in the intensity of MS peaks rather than to the presence or absence of any specific peaks. Recently, a more accurate prediction (78%) of the classification of *An. stephensi* mosquitoes, that were either uninfected or infected with *P. berghei*, was obtained for spectral patterns of the thorax using machine learning algorithms [78]. Although the same experimental model was used in both studies, the differences in the performance could be attributed to several factors, such as the experimental conditions of mosquito infections, the delay between infective blood feeding and specimen sacrifice, the mosquito body part used in MS analyses, and the method adopted for data analysis. So far, no studies reported the use of MALDI-TOF MS profiling to assess arboviruses in mosquito vectors. These preliminary results need to be further confirmed and tested, notably with mosquitoes collected in the field. For the determination of mosquito infection status by MALDI-TOF MS, standardization is stronglhy recommended in order to get comparable and reproducable results. A simultaneous identification of mosquito vector species and detection of their associated viral, bacterial, or parasitic pathogens would constitute a major improvement for entomological diagnosis and to assess the risk of mosquito-borne disease outbreaks.

### 8. Assessment of MALDI-TOF MS profiling to study the microbiota of mosquitoes

The term “microbiota” refers to a set of populations of microorganisms (viruses, protists, bacteria, fungi) in a particular place or time. During the past decade, interest has grown in the role of microbiota, especially gut microbiota of mosquito vectors, and in host-parasite interactions [83,84]. In the context of the present review, the term “microbiota” mainly refers to the commensal intestinal bacteria in mosquitoes. During the aquatic stage, mosquito larvae feed on organic detritus, including microorganisms, some of which become part of gut microflora in larval and adult stages [85]. Mosquito gut microbiota has been shown to be indispensable for larval development and survival of adult mosquitoes. Research on the various roles that gut microbiota play in mosquito biology – female fertility, adult longevity, immunity, formation of peritrophic matrix, nutrition, protection from insecticides due to insecticide-degrading gut bacteria – remain largely unknown [86,87]. Studies in this research area may reveal the mechanisms involved in insect immunity system and provide insight into the factors that influence vector capacity and competence as well as insecticide resistance [88]. This knowledge may in turn help designing more locally adapted strategies to control the propagation of various mosquito-borne pathogens, such as arboviruses and malaria parasites [87,89].

In one of the first studies that attempted to isolate and reveal the widest possible spectrum of aerobic and anaerobic bacteria present in the midgut of several mosquito species (*An. gambiae* s.l., *Ae. albopictus*, and *Culex quinquefasciatus*), the investigators proceed to a large-scale isolation of bacteria through microbial culturomics [90]. The species identification of both mosquitoes and their intestinal bacteria, as well as the bacteria found in breeding water for laboratory-reared mosquitoes, was confirmed by PCR sequencing and MALDI-TOF MS. Up to 16 bacterial species belonging to 12 genera were identified in the midgut of *An. gambiae*, 11 species from 8 genera were identified in *Ae. albopictus*, and finally 5 species from 5 genera were identified in *Cx. quinquefasciatus*. The authors confirmed that most of isolated bacteria belong to the phyla Proteobacteria and Firmicutes [90]. Although some strictly anaerobic bacteria cannot be isolated in cultures, the large-scale isolation technique of culturomics allowed the isolation of 17 additional bacterial species in mosquito midgut microbiota that had not been previously reported. Most of the bacteria present in the water were found in the midgut of adult mosquitoes, hence underlying the direct influence of larval breeding water in the microbiota composition of mosquitoes.

In subsequent studies, other authors found similar results using MALDI-TOF MS for bacterial identification in the midgut of *Ae. aegytpi*, *Ae. albopictus*, *Cx. quinquefasciatus, An. arabiensis*, and *An. funestus* that were either reared in the laboratory or caught in the field. The authors however reported some differences in the numbers and species of isolated bacteria due to environmental factors between laboratory-reared and field collected specimens [91,92]. In the study conducted by Tandina *et al*. [90], the majority of the isolated bacteria belonged to the phyla Proteobacteria and, to a lesser extent, Firmicutes. Microbial composition and diversity were not affected by sex, storing condition (fresh vs preservation), preservative mode, or storage period (up to 3 months) [92]. The authors concluded that MALDI-TOF MS is a valid tool for determining the composition of microbiota in mosquitoes using culturomic technique.

One of the promising strategies under consideration for vector control relies on the massive and regular release of *Ae. aegypti* adult male mosquitoes artificially infected with the bacteria *Wolbachia*. When *Wolbachia*-infected adult male mosquitos’ mate with wild females, the eggs do not hatch through cytoplasmic incompatibility [93]. *Wolbachia* bacteria invades most tissues in mosquito, including the salivary glands, midgut, muscle, and the nervous system, while shortening the life expectancy of the mosquito [94]. To evaluate the impact of this strategy, a monitoring of the presence of *Wolbachia* in *Ae. aegypti* mosquitoes after the release in the field is necessary. In this context, the MALDI-TOF MS coupled with artificial intelligence was assessed to distinct *Wolbachia*-infected from uninfected mosquitoes using the head and thorax [95]. This straegy gave a high accuracy rate of classification, comparable to that of quantitative PCR, and even superior to the loop-mediated isothermal amplification. The high throughput assays coupled with the low cost per sample could be particularly relevant to detect the presence of *Wolbachia* in mosquitoes in the frame work of large-scale field trials implemented worldwide.

### 9. Assessment of MALDI-TOF MS profiling to monitor insecticide resistance

Insecticide reistance in mosquitoes is considered by WHO as a serious threat to any vector control programme. So far, methods to detect insecticide resistance rely on biological, molecular and biochemical tools, each having strength and weakness [96]. Adequate surveillance of insecticide resistance requires repetive measurement of mosquito susceptibility to insecticides in multiple (sentinel) sites and as such, the number of samples to be tested can be extremely high [97]. Consequently, more adequate tools are needed to discriminate between susceptible and insecticide resistant mosquitoes, especially when large scale or nationwide monitoring programmes are implemented. Until recently there were no reports or publications relating to the use of MALDI-TOF MS profiling to identify insecticide resistance in insects. Our team then carried out the first study to assess the potential of MALDI TOF MS to distinguish between susceptible and pyrethroid resistant *Ae. aegypti* by comparing the protein signatures of legs and/or thoraxes of laboratory and field caught populations having different susceptibility to deltamethrin [98]. For this demonstration, a susceptible reference laboratory *Ae. aegypti* species (line BORA, French Polynesia) was compared to three inbreeds *Ae. aegypti* lines from French Guiana, with distinct deltamethrin resistance genotype/phenotype. Interestingly a peak at 4870 Da was found significantly more abundant in the high pyrethroid resistant *Ae. aegypti* population compared to the susceptible one’s coming from either the lab or French Guinana. Further analyses however failed to find a positive association between kdr resistant markers (V410L, V1016G/I, and F1534C) and the discriminant peak (i.e., 4870Da). Although these preliminary results are promising, further work is needed to characterise the peak of interest and to validate it as a marker of deltamethrin resistant in *Ae. aegypti* populations [98]. This work opens new perspectives of research in the field of insecticide resistance with the aim to facilitate the monitoring of insecticide resistance by national programmes.

### 10. Novel applications of MALDI-TOF MS: future perspectives

The MALDI-TOF MS profiling has demonstrated its interest for analyzing various mosquito life straits using specimens from different species and geographical origin. These classifications were generally done by spectral matching which could be insufficient for the detection of specific criteria. As previously mentioned, the association between artificial intelligence and the MALDI-TOF MS profiling have considerably improved the precision of identification as demonstrated with pathogen infections [78,95]. It is likely that machine learning algorithms will be applied and used extensively in the coming years, for the classification of MS spectra derived from mosquitoes, notably to distinct complex-or sibling-species, the microbiota composition and/or other key biological parameters.

In MALDI-TOF MS, the analysis of intact protein profiling (IPP) presents some limitations as previously observed with the determination of the blood feeding origin. Effectively, the digestion of blood protein impaired MS spectra matching with those of the reference database, reducing the host identification success when the processing time exceeds 24 hours post-blood feeding [43]. To extend the delay of host identification, the peptide mass mapping (PMM) or fingerprinting (PMF), consisting to analyse host-specific hemoglobin peptides using MALDI-TOF MS was assessed using bloodfed female of *Phlebotomus* spp. (vectors of *Leishmania* spp.) and laboratory reared *Culex* mosquitoes [99]. The principle is to digest the blood from arthropod abdomens by exogenous trypsin hence resulting in peptide fragments which serves as unique host signature in MALDI-TOF. As the half-life of hemoglobin peptides is longer than the respective whole proteins, PMM is less affected by blood degradation and these tryptic maps allowed conclusive host assignment up to 48 hours after the blood meal intake. Interestingly, the application of the PMF to distinguish closely-related *Culicoides* species strongly improved their classification [100]. The PMM approach appeared as a reliable alternative to study the blood source origin in mosquitoes and could be extended to other characters, especially for the identification of closely related species or for mosquito exuviae studies [72].

The MALDI-TOF MS profiling, in addition to protein investigations, has emerged as a potential alternative tool for the separation and detection of nucleic acids. Effectively, it has been employed to differentiate genotypes based on the mass of variant DNA sequences [101]. Among the studies related with mosquito life traits, three studies combining the detection and characterisation of nucleotidic changes by MALDI-TOF MS have been conducted. The first one addressed the detection and typing of dengue virus [102]. The principle was the comparison of RNA fragment profiles resulting from digestion by RNase T1 endoribonuclease of specific dengue virus PCR products, with *In silico* database of digestion patterns from dengue strains. This work showed the proof-of-concept for classifying hundreds of dengue virus down to the serotype and strain level. This work revealed the high potential of this strategy for an accurate dengue virus biotyping which could be extended to other arboviruses.

The two other works investigated the efficiency of MALDI-TOF MS to genotype single nucleotide polymorphism (SNP) sites in mosquito genes involved in insecticide resistance [103,104]. The principle consists to amplify gene containing SNP target and to extend probe juxtaposing the SNP site using ddNTPs. The base type of the target site is then determined by the mass of the extended probe by MALDI-TOF MS. The multiplexing of amplified target sites allows the simultaneous detection of multiple mutation sites. Five polymorphisms in the *acetylcholinesterase-1* gene, related to insecticide resistance in *Anopheles* spp. and *Culex* spp.. were analysed by Mao *et al* [103], whereas three mutations in the gene coding for the voltage-gate sodium channels, the main target of pyrethroid insecticides, generating 17 genotypes, from *Ae. albopictus* mosquitoes were investigated by Mu *et al* [104]. In both studies, the comparison of the multiplex PCR-mass spectrometry with conventional molecular methods indicated consistent genotyping results, confirming the accuracy of this approach. This technology allowed the rapid screening of several mutations at one single reaction per sample, decreasing the cost of the analysis to one-fifth [103]. Further application and development of this competitive strategy for rapid and reliable determination of other mosquito life traits based on SNPs, may be promising for surveillance programmes.

The MALDI mass spectrometry imaging (MALDI-MSI) that uses high-resolution images based on protein profiles represent another promising application of MALDI-TOF MS in medical entomology [105]. MALDI-MSI is a powerful tool that provides information on chemical composition, as well as the spatial distribution of molecules, from a sample sliced and loaded onto a glass slide. The images are created based on the mass-to-charge ratio of ions of interest measured by MALDI-TOF MS. In addition, MALDI-MSI has the advantages of being highly sensitive and, unlike other classical imaging methods, it can analyse from hundreds to thousands of molecules simultaneously in a single run, without labelling or altering the scanned tissue. This technology opens a way to spatial analysis of a range of analytes, including peptides, proteins, protein modifications, drugs and their metabolites or lipids. It offers then the mapping of specific molecules, in the arthropod specimens, like endosymbiote or pathogen proteins, the distribution of insecticides as well as protein repertory adaptation to environmental changes [106]. Until now, only one publication reported the application of MALDI-MSI to investigate the phospholipid composition, distribution, and localization in whole-body sections of the malaria vector, *An. stephensi* [107]. Such studies may represent the first step towards further understanding of parasite-host interaction, in particular lipid biochemistry underlying malaria infection, which in turn may reveal potential drug targets.

## Conclusion

The contribution of MALDI-TOF MS to biological sciences and medicine has been demonstrated for more than 20 years. Here we demonstrated that this technique has huge potential for application in medical entomology, especially for entomology surveillance and for the identification of various mosquito life traits. MALDI-TOF MS is a versatile and accurate tool that can be used to determine mosquito species, trophic preference, age and pathogen infections in mosquitoes. The establishment of species-specific MS protein signatures regardless of the mosquito development stages, from eggs to imago including exuviae, allows identification of mosquito species throughout their all life cycle. For adult mosquitoes, the combination of different body parts enhances the accuracy of identification, which could be decisive for identifying sibling species among complexes. This proteomic tool can be beneficial for surveillance programmes, as it showed to provide early and rapid identification of invasive vector species. Despite the numerous advantages of MS profiling, including its high cost-effectiveness, rapidity, and high throughput assays compatible with large-scale mosquito monitoring programmes, its use in entomology remains confidential. The lack of an international reference MS spectral database probably explains the limited interest for this technique although major progress has been made in this field. However, the continuous emergence and reemergence of mosquito-borne diseases worldwide underline the necessity to develop more sensitive technologies to improve the evaluation, surveillance, and prediction of epidemics. Integration of MALDI-TOF MS profiling as part of entomological surveillance could contribute to reduce the burden of mosquito borne diseases and guide decision making for vector control.

## List of abbreviations

BOLD: Barcode of Life Data Systems
COI: cytochrome c oxidase subunit I
COVID-2019: coronavirus disease 2019
ITS2: internal transcribed spacer
LAMP: loop-mediated isothermal amplification
MALDI-MSI: MALDI mass spectrometry imaging
MALDI-TOF MS: matrix-assisted laser desorption/ionization time-of-flight mass spectrometry
MS: mass spectrometry
NCBI: National Centre for Biotechnology Information
PMM: peptide mass mapping
PRISMA: Preferred Reporting Items for Systematic Reviews and Meta-Analyses
RT-PCR: reverse transcriptase-polymerase chain reaction
SNP: single nucleotide polymorphism

## Declarations

### Ethics approval and consent to participate

Not applicable.

### Consent for publication

Not applicable.

### Availability of data and materials

The complete set of search terms after selection of filters with respective results of articles retrieved per publication databases is provided in the Additional file S1.

### Authors’ contributions

Conceived and designed the experiments: LA and VC. Analyzed the data: LA, MMC, VC and RBH. Contributed reagents/materials/analysis tools: LA, MMC and RBH. Drafted the paper: LA, MMC and VC. Revised critically the paper: all authors.

## Acknowledgments

We thank the “Fondation Méditerranée Infection (FMI)” which offered personnel grant to MMC and RBH.

