## Additional_file_S1 for "MALDI-TOF MS profiling and its contribution to mosquito-borne diseases: a systematic review"

**Additional file S1**. Links of articles’ records from search of the databases.

| **Databases**  **(extraction date)** | **Search terms** | **Link of search*** |
| --- | --- | --- |
| **Pubmed (17.02.23)** | Maldi AND Mosquito | https://pubmed.ncbi.nlm.nih.gov/?term=Maldi+AND+Mosquito&filter=dates.2003%2F1%2F1-2023%2F2%2F17&filter=lang.english&filter=other.excludepreprints&size=200 |
| **Web of Science (17.02.23)** | Maldi AND Mosquito | https://www.webofscience.com/wos/woscc/summary/f5303ea2-2b86-4515-b37b-81b67d97ee85-715e2eab/relevance/1 |
| **Science Direct (16.02.23)** | **1. (Maldi OR "mass spectrometry") AND (Mosquito) AND (surveillance OR monitoring OR management)** | |
|  | Maldi AND "mosquito surveillance" | https://www.sciencedirect.com/search?qs=Maldi%20AND%20%22mosquito%20surveillance%22&date=2003-2023&lastSelectedFacet=articleTypes&articleTypes=REV |
|  | Maldi AND "mosquito monitoring" | https://www.sciencedirect.com/search?qs=Maldi%20AND%20%22mosquito%20monitoring%22&years=2016&lastSelectedFacet=years |
|  | "mass spectrometry" AND "mosquito surveillance" | https://www.sciencedirect.com/search?qs=%22mass%20spectrometry%22%20AND%20%22mosquito%20surveillance%22&date=2003-2023&articleTypes=REV%2CFLA&lastSelectedFacet=articleTypes |
|  | "mass spectrometry" AND "mosquito monitoring" | https://www.sciencedirect.com/search?qs=%22mass%20spectrometry%22%20AND%20%22mosquito%20monitoring%22&date=2003-2023&lastSelectedFacet=articleTypes&articleTypes=FLA |
|  | "mass spectrometry" AND "mosquito management" | https://www.sciencedirect.com/search?qs=%22mass%20spectrometry%22%20AND%20%22mosquito%20management%22&date=2003-2023&articleTypes=FLA%2CREV&lastSelectedFacet=articleTypes |
|  | **2. (Maldi OR "mass spectrometry") AND (Mosquito) AND ("Mosquito Identification")** | |
|  | Maldi AND "mosquito identification" | https://www.sciencedirect.com/search?qs=Maldi%20AND%20%22mosquito%20identification%22&years=2022&lastSelectedFacet=years |
|  | "mass spectrometry" AND "mosquito identification" | https://www.sciencedirect.com/search?qs=%22mass%20spectrometry%22%20AND%20%22mosquito%20identification%22 |
|  | **3. (Maldi OR "mass spectrometry") AND (Mosquito) AND (Longevity OR Age OR oviposition)** | |
|  | Maldi AND "mosquito longevity" | https://www.sciencedirect.com/search?qs=Maldi%20AND%20%22mosquito%20longevity%22 |
|  | Maldi AND "mosquito age" | https://www.sciencedirect.com/search?qs=Maldi%20AND%20%22mosquito%20age%22&date=2003-2023&articleTypes=REV%2CFLA&lastSelectedFacet=articleTypes |
|  | Maldi AND "mosquito oviposition" | https://www.sciencedirect.com/search?qs=Maldi%20AND%20%22mosquito%20oviposition%22 |
|  | "mass spectrometry" AND "mosquito longevity" | https://www.sciencedirect.com/search?qs=%22mass%20spectrometry%22%20AND%20%22mosquito%20longevity%22&date=2003-  2023&articleTypes=FLA&lastSelectedFacet=articleTypes |
|  | "mass spectrometry" AND "mosquito age" | https://www.sciencedirect.com/search?qs=%22mass%20spectrometry%22%20AND%20%22mosquito%20age%22&date=2003-2023&articleTypes=REV%2CFLA&lastSelectedFacet=articleTypes |
|  | "mass spectrometry" AND "mosquito oviposition" | https://www.sciencedirect.com/search?qs=%22mass%20spectrometry%22%20AND%20%22mosquito%20oviposition%22&date=2003-2023&articleTypes=REV%2CFLA&lastSelectedFacet=articleTypes |
|  | **4. (Maldi OR "mass spectrometry") AND (Mosquito) AND (Feed OR Blood OR engorged or "Feed behavior")** | |
|  | Maldi AND "mosquito feed" | https://www.sciencedirect.com/search?qs=Maldi%20AND%20%22mosquito%20feed%22&date=2003-2023&articleTypes=FLA&lastSelectedFacet=articleTypes |
|  | Maldi AND "mosquito blood" | https://www.sciencedirect.com/search?qs=Maldi%20AND%20%22mosquito%20blood%22&date=2003-2023&articleTypes=REV%2CFLA&lastSelectedFacet=articleTypes |
|  | Maldi AND "mosquito engorged" | https://www.sciencedirect.com/search?qs=Maldi%20AND%20%22mosquito%20engorged%22 |
|  | Maldi AND "mosquito feed behavior" | https://www.sciencedirect.com/search?qs=Maldi%20AND%20%22mosquito%20feed%20behavior%22 |
|  | "mass spectrometry" AND "mosquito feed" | https://www.sciencedirect.com/search?qs=%22mass%20spectrometry%22%20AND%20%22mosquito%20feed%22&date=2003-2023&articleTypes=REV%2CFLA&lastSelectedFacet=articleTypes |
|  | "mass spectrometry" AND "mosquito blood" | https://www.sciencedirect.com/search?qs=%22mass%20spectrometry%22%20AND%20%22mosquito%20blood%22&date=2003-2023&articleTypes=REV%2CFLA&lastSelectedFacet=articleTypes |
|  | "mass spectrometry" AND "mosquito engorged" | https://www.sciencedirect.com/search?qs=%22mass%20spectrometry%22%20AND%20%22mosquito%20engorged%22 |
|  | "mass spectrometry" AND "mosquito feed behavior" | https://www.sciencedirect.com/search?qs=%22mass%20spectrometry%22%20AND%20%22mosquito%20feed%20behavior%22 |
|  | **5. (Maldi OR "mass spectrometry") AND (Mosquito) AND (origin OR geographic)** | |
|  | Maldi AND "mosquito origin" | https://www.sciencedirect.com/search?qs=Maldi%20AND%20%22mosquito%20origin%22&date=2003-2023 |
|  | Maldi AND "mosquito geographic" | https://www.sciencedirect.com/search?qs=Maldi%20AND%20%22mosquito%20geographic%22 |
|  | "mass spectrometry" AND "mosquito origin" | https://www.sciencedirect.com/search?qs=%22mass%20spectrometry%22%20AND%20%22mosquito%20origin%22&years=2008%2C2009%2C2020&lastSelectedFacet=years |
|  | "mass spectrometry" AND "mosquito geographic" | https://www.sciencedirect.com/search?qs=%22mass%20spectrometry%22%20AND%20%22mosquito%20%20geographic%22 |
|  | **6. (Maldi OR "mass spectrometry") AND (Mosquito) AND ( "Mosquito disease" OR "Mosquito-borne-disease" OR MBD OR Parasite OR Arbovirus)** | |
|  | Maldi AND "Mosquito disease" | https://www.sciencedirect.com/search?qs=Maldi%20AND%20%22Mosquito%20disease%22&articleTypes=REV&lastSelectedFacet=articleTypes |
|  | Maldi AND "Mosquito-borne-disease" | https://www.sciencedirect.com/search?qs=Maldi%20AND%20%22Mosquito-borne-disease%22&date=2003-2023&articleTypes=REV%2CFLA&lastSelectedFacet=articleTypes |
|  | Maldi AND "Mosquito Parasite" | https://www.sciencedirect.com/search?qs=Maldi%20AND%20%22Mosquito%20Parasite%22&date=2003-2023&articleTypes=REV%2CFLA&lastSelectedFacet=articleTypes |
|  | Maldi AND "Mosquito Arbovirus" | https://www.sciencedirect.com/search?qs=Maldi%20AND%20%22Mosquito%20Arbovirus%22 |
|  | "mass spectrometry" AND "Mosquito disease" | https://www.sciencedirect.com/search?qs=%22mass%20spectrometry%22%20AND%20%22Mosquito%20disease%22&date=2003-2023&articleTypes=REV%2CFLA&lastSelectedFacet=articleTypes |
|  | "mass spectrometry" AND "Mosquito-borne-disease" | https://www.sciencedirect.com/search?qs=%22mass%20spectrometry%22%20AND%20%22Mosquito-borne-disease%22&date=2003-2023&articleTypes=REV%2CFLA&lastSelectedFacet=articleTypes |
|  | "mass spectrometry" AND "Mosquito Parasite" | https://www.sciencedirect.com/search?date=2003-2023&qs=%22mass%20spectrometry%22%20AND%20%20%22Mosquito%20Parasite%22&articleTypes=REV%2CFLA&lastSelectedFacet=articleTypes |
|  | "mass spectrometry" AND "Mosquito Arbovirus" | https://www.sciencedirect.com/search?qs=%22mass%20spectrometry%22%20AND%20%22Mosquito%20Arbovirus%22 |
|  | **7. (Maldi OR "mass spectrometry") AND (Mosquito) AND (Bacteria OR Wolbachia)** | |
|  | Maldi AND "Mosquito Bacteria " | https://www.sciencedirect.com/search?date=2003-2013&qs=Maldi%20AND%20%22Mosquito%20Bacteria%22&articleTypes=FLA&lastSelectedFacet=articleTypes |
|  | Maldi AND "Mosquito Wolbachia" | https://www.sciencedirect.com/search?qs=Maldi%20AND%20%22Mosquito%20Wolbachia%22 |
|  | "mass spectrometry" AND "Mosquito Wolbachia" | https://www.sciencedirect.com/search?date=2003-2023&qs=%22mass%20spectrometry%22%20AND%20%22Mosquito%20Wolbachia%22 |
|  | "mass spectrometry" AND "Mosquito Bacteria " | https://www.sciencedirect.com/search?qs=%22mass%20spectrometry%22%20AND%20%22Mosquito%20Bacteria%20%22 |
|  | **8. (Maldi OR "mass spectrometry") AND (Mosquito) AND (“Insecticide resistance" OR "Insecticide susceptibility")** | |
|  | Maldi AND "Mosquito" AND “Insecticide resistance" | https://www.sciencedirect.com/search?qs=Maldi%20AND%20%22Mosquito%22%20AND%20%E2%80%9CInsecticide%20resistance%22&date=2003-2023&articleTypes=REV%2CFLA&lastSelectedFacet=articleTypes |
|  | Maldi AND "Mosquito" AND “Insecticide susceptibility" | https://www.sciencedirect.com/search?qs=Maldi%20AND%20%20%22Mosquito%22%20AND%20%E2%80%9CInsecticide%20susceptibility%22&date=2003-2023&articleTypes=FLA&lastSelectedFacet=articleTypes |

*Search results after selection of filters (*i.e*. year frame, type of article and language).
